# Universal-Bac^3^Gel^®^: a 3D Biofilm-Relevant Matrix that Supports *In Vitro* Growth and Biofilm Formation of ESKAPE Pathogens

**DOI:** 10.64898/2025.12.16.694620

**Authors:** Emanuela Peluso, Sebastião Van Uden, Sonja Visentin, Paola Petrini, Daniela Peneda Pacheco, Livia Visai

## Abstract

Human microbiota is increasingly considered to shape health and disease, drawing interest of pharma and biotech industries in advanced models of *in vitro* human microbiome to streamline drug development. In this context, Universal-Bac^3^Gel^®^ represents a new generation of 3D biomaterials designed to mimic the properties of human mucus and biofilm features, including micro-gradients that replicate the heterogeneous environments colonized by microorganisms in the human body. To evaluate the suitability of Universal-Bac^3^Gel^®^ for studying clinically relevant species in antimicrobial resistance, the so-called ESKAPE pathogens (*Enterococcus faecium, Staphylococcus aureus, Klebsiella pneumoniae, Acinetobacter baumannii, Pseudomonas aeruginosa, and Enterobacter cloacae*) were cultured within this 3D environment. Bacterial growth was monitored at 24- and 48-hours post-inoculation via spot plating, while viability, distribution and organization were assessed using confocal laser scanning microscopy. All ESKAPE strains successfully grew throughout the structure of Universal-Bac^3^Gel^®^. Distinct 3D biofilm architectures were observed across species, ranging from diffuse colonization to compact microcolony formation, in agreement with species-specific biofilm patterns. Notably, the platform’s ready-to-use 96-well format allowed direct comparison of these high-priority pathogens under identical conditions, highlighting species-specific biofilm traits that would be difficult to discern in traditional 2D or animal models. This work highlights the versatility of Universal-Bac^3^Gel^®^ as a platform for studying pathogen colonization under a biofilm-relevant environment and underscores its potential for translational applications, including antimicrobial drug testing, personalized medicine, and reduction of animal model use in preclinical research.

## 1. Introduction

The human resident bacterial flora, i.e., all microbial populations that establish a symbiotic relationship with the tissues of the human organism, is collectively known as the microbiota. Current quantitative analyses indicate that the total number of microbial and human cells in the body is approximately equal, with a near 1:1 ratio, although this estimate varies depending on host-specific factors such as body size, diet, and microbiome composition (Sender et al., 2016). While all human organisms are anatomically similar, multiple factors influence the development of bacterial populations, determining the specificity of the microbiota for each individual.

Despite remarkable advances in sequencing technologies and microbial ecology, a substantial fraction of microorganisms in both natural and human-associated environments remains uncultured, limiting our understanding of their functional roles in health and disease. In nature, bacteria predominantly grow as biofilms-structured, surface-associated communities embedded in a self-produced extracellular matrix. It is estimated that roughly 99% of bacteria on Earth exist in biofilms, while only about 1% persist in a free-floating, planktonic state (Flemming & Wuertz, 2019). Nevertheless, most *in vitro* studies still rely on planktonic cultures, which fail to reproduce the complex structural and physiological characteristics of bacteria growing *in vivo*.

To address these limitations, innovative cultivation strategies have emerged over the past decade, including growth in natural habitats and the use of modified media that enable a broader recovery of microbial diversity. However, conventional two-dimensional (2D) models for studying bacterial infections remain limited in their ability to accurately reproduce the *in vivo* environment, as they lack pathogen-specific cell types, intercellular signaling, and biomechanical cues (Fasciano et al., 2019); (Bermudez-Brito et al., 2013); (Shi et al., 2019).

These shortcomings become particularly critical when investigating infections at mucosal surfaces, where the biochemical and physical microenvironment plays a key regulatory role. Mucosal surfaces constitute the primary ecological niche for microbial colonization and infection. They are coated with mucus layers composed of mucins, lipids, and other proteins such as immunoglobulins, which serve both as a physical barrier and as a dynamic biochemical habitat for bacteria (Cone, 2009). The viscoelastic and heterogeneous nature of mucus creates spatial gradients of oxygen, nutrients, and antimicrobial peptides that strongly influence microbial behavior and biofilm development (Lieleg & Ribbeck, 2011). These features are rarely reproduced in traditional *in vitro* systems, underscoring the need for advanced models that more faithfully mimic the biophysical properties of mucus, as well as the unified mucus and biofilm niche - mucofilm (Mortensen et al., 2025).

Recent technological advances highlight the promise of three-dimensional (3D) organotypic systems and organ-on-a-chip platforms that replicate physiological and pathological tissue functions with greater fidelity, providing improved tools to study host-microbe interactions and infection dynamics (Rikken et al., 2023); (Nasiri et al., 2024). At the same time, the escalating global challenge of antimicrobial resistance highlights the importance of physiologically relevant systems to investigate pathogen adaptation, persistence, and treatment response (Santos & Coelho, 2025); (Higazy et al., 2024).

In particular, surveillance data have identified a group of nosocomial bacteria - collectively known as the ESKAPE pathogens - that are responsible for the majority of hospital-acquired infections worldwide. This group includes both Gram-positive (**Figure 1** purple) and Gram-negative species (**Figure 1** blue), notorious for multidrug resistance and their ability to “escape” the effects of antibiotics (Santajit & Indrawattana, 2016); (Bush & Jacoby, 2010). These organisms represent leading causes of severe infections, particularly among critically ill or immuno-compromised patients (Ram et al., 2010).

**Figure 1.**
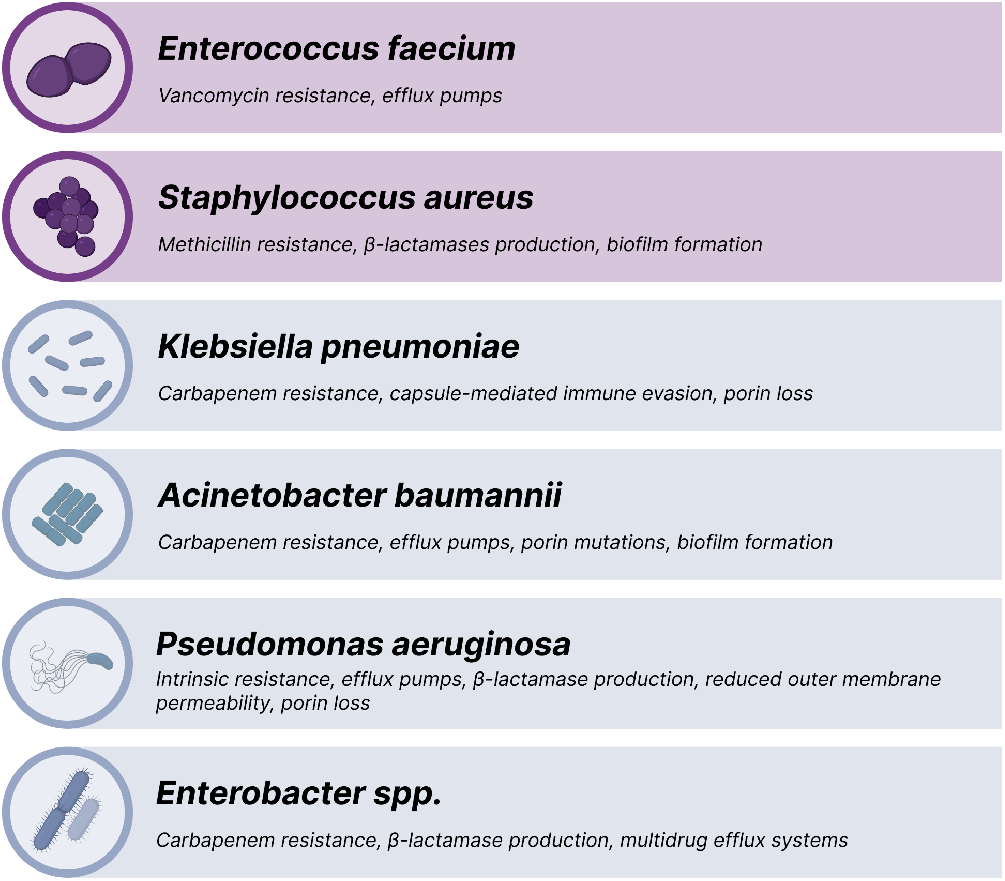
Overview of the ESKAPE pathogens, divided into Gram-positive (purple) and Gram-negative (blue) groups, and their major antimicrobial resistance mechanisms. *The ESKAPE group comprises six clinically significant species -Enterococcus faecium* (Santajit & Indrawattana, 2016) (Munita & Arias, 2016); *Staphylococcus aureus* (Pendleton et al., 2013) (Munita & Arias, 2016) *Klebsiella pneumoniae* (Pendleton et al., 2013) (Santajit & Indrawattana, 2016); *Acinetobacter baumannii* (Pendleton et al., 2013) (Santajit & Indrawattana, 2016), *Pseudomonas aeruginosa* (Munita & Arias, 2016) (Boucher et al., 2009) *and Enterobacter spp*. (Santajit & Indrawattana, 2016). *Each exhibits distinct resistance strategies, including β-lactamase production, efflux pumps, porin modifications, and biofilm formation*.

In this context, Universal-Bac^3^Gel® represents an innovative 3D biomaterial designed to mimic the structural, biochemical, and rheological properties of human mucus and biofilm matrices. The hydrogel reproduces essential features such as oxygen and nutrient gradients, diffusion limitations, and viscoelastic behavior (storage modulus G′ > G″), which collectively create a physiologically relevant environment for microbial colonization (Pacheco et al., 2023). By capturing these microenvironmental cues, Universal-Bac^3^Gel® facilitates studies in pharmacokinetics, antimicrobial testing, microbiota reconstruction, and microbial ecology (Butnarasu et al., 2022) (Vargas et al., 2025). Several *in vitro* and *ex vivo* systems have been developed to mimic mucus-like environments and biofilm formation (Crabbé et al., 2019); (Harrison & Diggle, 2016). Among them, artificial sputum medium (ASM) and its modified versions (ASM_mod) are widely used to replicate cystic fibrosis (CF) airway mucus (Aiyer & Manos, 2022). Compared to other established models, such as ASM, AirGels, or organotypic tissue co-cultures (Möckel et al., 2022); (Phogat et al., 2023), Universal-Bac^3^Gel® offers distinct advantages. Its ready-to-use format in 96-well plates ensures reproducibility, scalability, and compatibility with high-throughput assays, making it suitable for both fundamental microbiology research and drug discovery. Importantly, the presence of biofilm-relevant physicochemical gradients allows the model to better reproduce the complex spatial and metabolic heterogeneity observed *in vivo*, resulting in more predictive results. The translational relevance of such physiologically accurate *in vitro* platforms is increasingly recognized. They offer the potential to evaluate novel antimicrobials or anti-biofilm strategies prior to animal testing or clinical trials. Given the current shortage of innovative antibiotics, with only 15 of 90 candidates considered truly novel and few targeting critical ESKAPE pathogens (Global Antibiotic Resistance Surveillance Report 2025 WHO Global Antimicrobial Resistance and Use Surveillance System (GLASS), a biofilm-relevant matrix like Universal-Bac^3^Gel^®^ can serve as a robust screening platform. By replicating infection-*like* microenvironments, it may uncover antimicrobial activities or tolerance mechanisms that conventional planktonic assays fail to detect, thereby improving the selection of promising drug candidates. Ultimately, this could streamline R&D by focusing resources on compounds effective in biofilm/mucus contexts, where many antibiotics notoriously fail. Understanding the intricate interactions between microbes, their environment, and host tissues remains essential for developing physiologically relevant *in vitro, ex vivo*, and *in vivo* models (Gilbert et al., 2018); (Barrila et al., 2018). This study aims to address this gap by developing a model that faithfully recapitulates the physicochemical and biological properties of mucosal environments to investigate microbial colonization and resistance mechanisms.

## 2. Materials and Methods

### 2.1 Bac^3^Gel^®^ technology

By replicating key properties of the biofilm environment, Bac^3^Gel^®^ (https://www.bac3gel.com/) provides an advanced 3D material exhibiting micro-gradients for the culture of microorganisms. In this study, Universal-Bac^3^Gel^®^ was exploited as a general-use 3D support for bacteria colonization and growth, which is provided in a standard 96-well flat-bottomed sterile microplate, as ready to use. Universal-Bac^3^Gel^®^ composition is formulated to reproduce the biofilm stratified environment, creating a diffusion-limited environment with oxygen and nutrient gradients similar (Pacheco et al., 2023); (Vargas et al., 2025). A detailed step-by-step protocol for culturing single strains in Universal-Bac^3^Gel® is available in Appendix 1 (Link A1).

### 2.2 Rheological Characterization

The viscoelastic properties of Universal-Bac^3^Gel^®^ produced in different media (7.07 mg mL^−1^ NaCl, MHB, and LB) were evaluated using an Anton Paar MCR501 Rheometer (Austria) with a 25 mm diameter plate geometry (serial number 52530/19910) at 25 °C. The linear viscoelastic region (LVR) was determined through strain sweep analyses employing a logarithmic ramp strain varying from 0.1% to 1000% at a frequency of 1 Hz. Oscillatory frequency sweeps were further performed to evaluate both storage, *G’*, and dissipative, *G″*, moduli, at 0.5% (at strain amplitudes within the linear regime) with frequencies changing logarithmically in the 0.1-20 Hz range.

### 2.3 Oxygen Tension Measurements

O_2_ tension of sterile Universal-Bac^3^Gel^®^ was measured using a Clark-type O_2_ sensor (OX-25; Unisense, Aarhus N, Denmark), connected to the Unisense microsensor multimeter S/N 8678 (Unisense, Denmark), a high sensitivity pico-ampere four-channel amplifier. Before each measurement, the reference anode and the guard cathode were polarized overnight and calibrated with either water saturated with air or with an anoxic solution of 2% (w/w) sodium hydrosulfite. Once the calibration was carefully made, a low melting point agarose consisting of 2% (w/v) agarose in 0.071% (w/v) NaCl (same concentration as in Universal-Bac^3^Gel^®^) was cast into a petri dish. Universal-Bac^3^Gel^®^ were placed over the agarose layer. Microsensors with a tip diameter of 50 µm (OX-50) were positioned at the air-Bac^3^Gel^®^interface (“depth zero”) using a motorized micromanipulator (Unisense, Denmark). Measurements were performed at the center of the hydrogels starting at their surface (0 mm) through their thickness, every 100 µm in triplicate, until the tip completely penetrated the whole structure. The maximum depth reached by the tip, termed end depth, was 3000 µm. The Unisense software SensorTrace automatically converts the signal from partial pressure (O_2_ tension) to the equivalent O_2_ concentration in µmol L^−1^.

### 2.4 Bacterial Strains and Culture Conditions

The microorganisms used were *Enterococcus faecium* (*E. faecium*), *Staphylococcus aureus* ATCC 25923 (*S. aureus*), *Pseudomonas aeruginosa* PAO1 (*P. aeruginosa*), and *Enterobacter cloacae* (*E. cloacae*) kindly supplied by Prof. Livia Visai (Department of Molecular Medicine, University of Pavia, Italy) and stored at -80°C; *Klebsiella pneumoniae* ATCC 13883 (*K. pneumoniae*), *Acinetobacter baumannii* LMG 1025 (*A. baumannii*) were kindly provided by Doctor Ângelo Filipe Santos Luis (Health Sciences Research Center, Beira Interior University) and stored at -80°C. Bacteria were grown in their appropriate medium overnight, under aerobic conditions, at 37°C using a shaker incubator (VDRL Stirrer 711/CT, Asal S.r.l., Milan, Italy). Brain Heart Infusion (BHI, Sigma-Aldrich, 53286, Lot#BCCD5575) was used for *K. pneumoniae, P. aeruginosa, S. aureus, E. faecium* and *E. cloacae*. Mueller Hinton (MH, Sigma-Aldrich, 70192, Lot# BCCC5707) was used to grow *A. baumannii* LMG 1025. *E. coli* were grown in Luria Bertani broth (ForMedium™, Hunstanton, Norfolk, UK). The inocula were diluted to a final concentration of 10^4^ bacteria/mL as determined by comparing the optical density (OD_600_) of the sample with a standard curve relating OD_600_ to cell number. (Restivo et al., 2024). Studies focused on the ESKAPE panel of six nosocomial pathogens (which exhibit multidrug resistance and high virulence (Mulani et al., 2019) given their priority status in new antibiotic development efforts. To evaluate the growth in both conditions, 100 µl of bacterial suspension was incubated overnight in Universal-Bac^3^Gel^®^, at 37°C. Inoculations were carried out for 24 and 48 hours. Planktonic cultures were conducted as controls of the experiment, while sterile BHI, MH, and Luria Bertani broth, as well as sterile Universal-Bac^3^Gel were used as controls of sterility.

### 2.5 Spot plating and colony-forming unit (CFU) counting

The spot plating method was selected as a substitute for the conventional plating method to avoid time-consuming steps, reduce material costs, and obtain high-throughput data (Zeden & Gröndling, 2023). After 24 and 48 hours of colonization, bacteria grown in Universal-Bac^3^Gel^®^ were determined through CFU plating. Before performing the dilutions, the Universal-Bac^3^Gel® was dissolved using Bac^3^Gel^®^ dissolution medium (50 mM sodium citrate) to retrieve bacteria grown within the hydrogel structure. Serial dilutions were performed using 96-well culture plates, with each well containing 90 µL of 0.9% NaCl, to which 10 µL of the bacterial suspension was added. After thorough pipetting in each well, several dilutions were made for the same starting sample. Using a calibrated micropipette, 10 μl aliquots from selected four dilutions were applied on top of the dedicated agar plate. The plates were left in the laminar air-flow chamber for the droplets to dry off (approx. 8–10 min), after which they were incubated overnight at 37°C to allow colony formation. Planktonic cultures were conducted as controls. For this, the same steps were followed, except for the dissolving agent. For the sterility controls, 10 µL of culture medium was taken from the 96-well plate used in the experiment and directly spotted onto an agar plate, as well as 10 µL of dissolved Universal-Bac^3^Gel^®^. Each experiment was performed in triplicate at two different times. After overnight incubation, CFU counting was performed manually by counting the individual colonies (dots) present on the agar plate.

### 2.6 Confocal laser scanning microscope analysis

Bacteria were grown overnight in Universal-Bac^3^Gel^®^ as previously described. After 24 and 48 hours of inoculation in Universal-Bac^3^Gel^®^, the supernatant was carefully removed, and the hydrogels were washed twice with Phosphate-Buffered Saline (PBS). Bacterial viability in Universal-Bac^3^Gel^®^ was assessed using the LIVE/DEAD® BacLight™ Viability Kit (Molecular Probes, Eugene, OR, USA). For staining, SYTO9^®^ was incubated for 10 minutes, followed by PI for 30 seconds. Bacteria were then observed using a Leica TCS SP8 DLS Confocal Laser Scanning Microscope (CLSM) (Leica, Wetzlar, Germany). Viable bacteria were visualized with a 25× multi-immersion phase contrast objective. The orthogonal projections and 3D reconstructions were acquired from different (n=5) regions of interest. Scale bars were generated using the LAS X software. The fluorescence intensity of the red and green channels of the 5 acquired images was evaluated using FIJI software. (Schindelin et al., 2012). This qualitative imaging complements the quantitative CFU data, providing insight into the 3D organization and viability of the bacterial communities inside the hydrogel.

### 2.7 Statistical analysis

All the statistical calculations were carried out using GraphPad Prism 9 (GraphPad Inc., San Diego, CA, USA). Statistical analysis was performed using two-sided t-test (significance level of p < 0.05). In addition, Two-Way ANOVA (or Mixed-Model), followed by Bonferroni’s multiple comparisons test was performed.

## 3. Results

### 3.1 Characteristics of Universal-Bac^3^Gel^®^

The hydrogel is a transparent, self-supporting cylindrical matrix compatible with standard multiwell formats (**Figure 2A**). Oxygen profiling along the gel depth revealed a stable gradient from approximately 290 µmol L^-1^ at the surface to ∼195 µmolL^-1^ near 3 mm depth (**Figure 2B**). Rheological characterization demonstrated that Universal-Bac^3^Gel® possesses viscoelastic properties consistent with extracellular polymeric substance (EPS)-rich matrices with a storage modulus (G′) higher than the loss modulus (G″) (**Figure 2C-D**), indicating solid-*like* behavior essential for maintaining biofilm structure under shear or mechanical stress. The moduli remained stable across gels hydrated with NaCl, Mueller-Hinton, or Luria-Bertani media, confirming mechanical robustness and reproducibility under standard microbial culture conditions.

**Figure 2.**
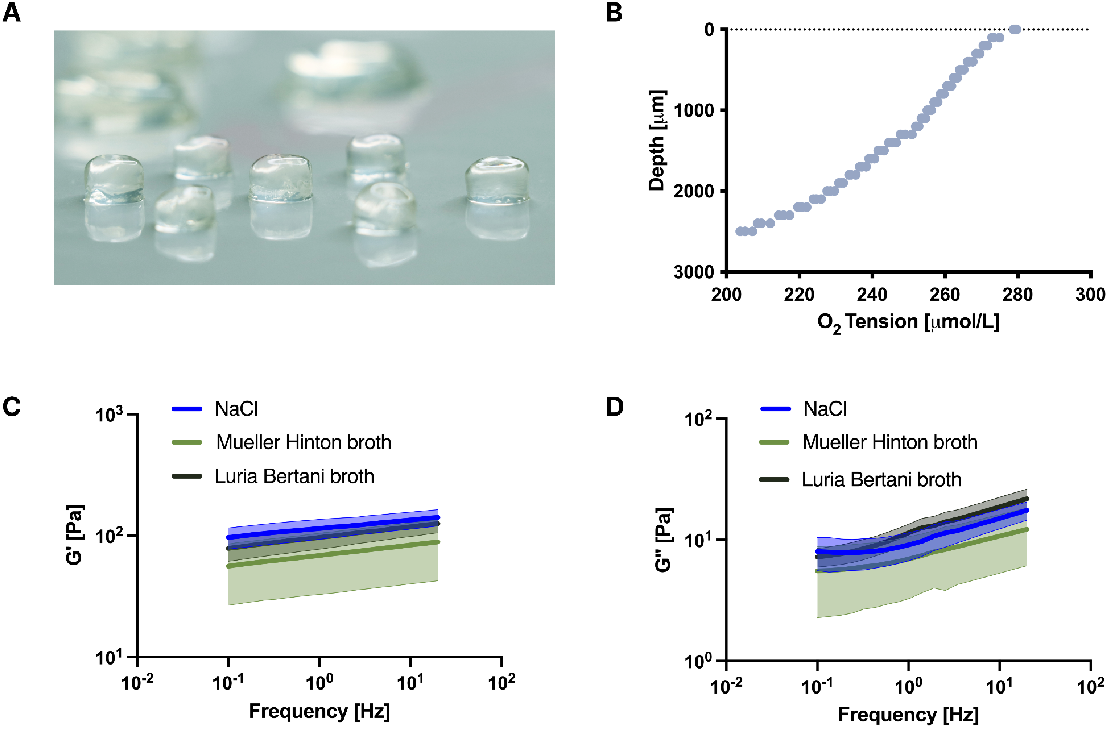
Characterization of Universal-Bac^3^Gel^®^ as a biofilm-relevant matrix. (**A**) Macroscopic image depicting the cylindrical-shape of Bac^3^Gel^®^; (**B**) Heterogeneous distribution of oxygen tension along Bac^3^Gel^®^ depth displaying a diffusion-driven gradient; (**C-D**) Frequency-dependent storage (G′) and loss (G″) moduli measured for gels hydrated with NaCl, Mueller-Hinton, or Luria-Bertani media. The area around the lines in **C** and **D** represents the standard deviation.

### 3.2 Colonization and biofilm formation by Gram-positive ESKAPE pathogens in Universal-Bac^3^**Gel**^**®**^

The two Gram-positive bacteria, *Enterococcus faecium* (Kim et al., 2021) and *Staphylococcus aureus* (Olsen et al., 2012), are major contributors to hospital-acquired infections and are known for their ability to form robust and persistent biofilms. To investigate the colonization potential of Gram-positive ESKAPE pathogens in a 3D biofilm-relevant environment, both strains were inoculated on top of Universal-Bac^3^Gel^®^ and their permeation and colonization were monitored over 24 and 48 hours. As shown in the CFU quantification, both *E. faecium* (**Figure 3A**) and *S. aureus* (**Figure 3D**) demonstrated high viability within the hydrogel, reaching concentrations ranging from 10^8^ to 10^15^ CFU/mL across both time points. These levels were statistically equivalent (p > 0.05) to planktonic culture controls (white bars), indicating that the matrix does not hinder bacterial growth and instead supports sustained proliferation over time.

**Figure 3.**
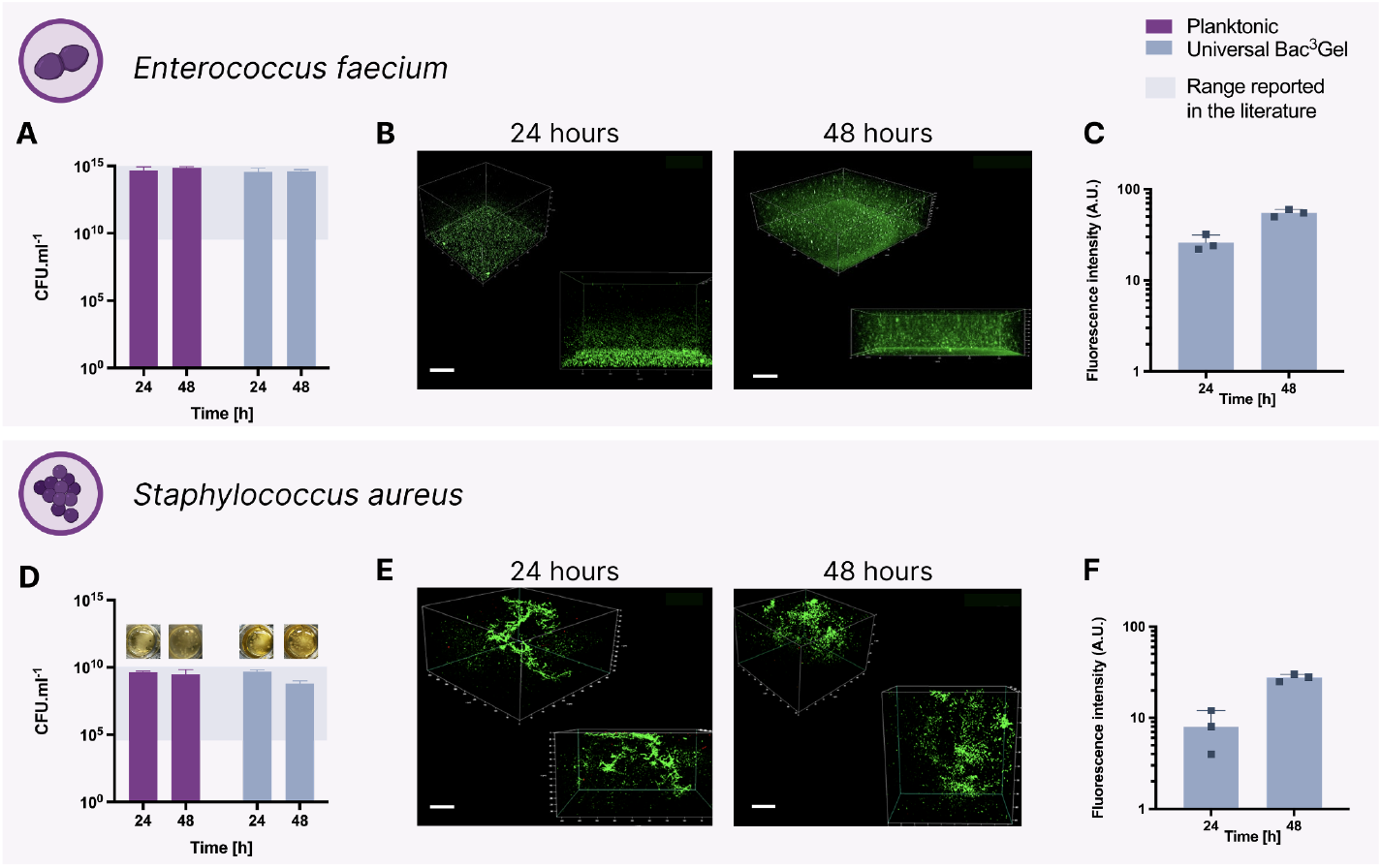
Growth and biofilm formation of Gram-positive ESKAPE pathogens in Universal-Bac^3^Gel^®^. (**A, D**) CFU counts of E. faecium and S. aureus at 24 h and 48 h in Universal-Bac^3^Gel^®^ (purple) compared with planktonic cultures (blue). The shaded grey area indicates the CFU/mL range reported for infected mucosa. Data are mean ± SD (n = 3); p < 0.05 for all comparisons relative to 0 h inoculum; no significant difference between Universal-Bac^3^Gel^®^ and planktonic were detected at each time point. Photographs were taken only for the commercially available reference strains and not for clinical isolates. (**B, E**) Representative CLSM images showing live/dead staining in Universal-Bac^3^Gel^®^. Scale bar: 50 µm. (**C, F**) Quantification of green fluorescence intensity corresponding to viable biofilm biomass) in 3D image stacks, illustrating total biofilm volume per field.

Microscopy-based qualitative analysis revealed distinct 3D spatial organization for the two Gram-positive strains (**Figure 3B, 3E**). *E. faecium* exhibited a relatively homogeneous distribution of green fluorescence throughout Bac^3^Gel^®^, forming a dense but dispersed biofilm (**Figure 3B**). The structure appeared compact, without large aggregates, suggesting a uniform colonization pattern reminiscent of a lawn-like biofilm. *S. aureus*, in contrast, displayed a markedly different architecture, forming dense and spatially localized aggregates typical of microcolony-based biofilm structures (**Figure 3E**). These clusters are consistent with the known ability of *S. aureus* to form mature and structured biofilms with high biomass density (Schilcher & Horswill, 2020).

Live/dead staining indicated predominantly live (green) cells in both cases, with only sparse dead (red) cells visible, suggesting the bacteria remained highly viable in the matrix. Quantification of fluorescence intensity in 3D reconstructions (**Figure 3C, 3F**) provided a measure of biofilm biomass. *E. faecium* (**Figure 3C**) displayed significantly brighter and more localized fluorescent regions, indicating higher biofilm density and possibly greater extracellular matrix production. In contrast, *S. aureus* (**Figure 3F**) showed a more moderate fluorescent signal that was more heterogeneously distributed, consistent with its fewer but larger clumps.

### 3.3 Colonization and biofilm formation by gram-negative ESKAPE pathogens in Universal-Bac3Gel®

The Gram-negative ESKAPE pathogens *Klebsiella pneumoniae, Acinetobacter baumannii, Pseudomonas aeruginosa*, and *Enterobacter cloacae* also represent critical agents of nosocomial infections and are characterized by their intrinsic and acquired resistance mechanisms, as well as their capacity to form complex biofilm communities (Santajit & Indrawattana, 2016). We evaluated their colonization potential in the 3D biofilm-relevant environment Universal-Bac^3^Gel^®^ with cultures at 24 and 48 hours. CFU analysis revealed that all strains remained viable within the hydrogel, maintaining high concentrations on the order of 10^6^-10^10^ CFU/mL across both time points (**Figure 4A, D, G, J**), similar to planktonic conditions. In most cases, CFU counts in Universal-Bac^3^Gel® were comparable to planktonic controls. Notably,*K. pneumoniae* and *E. cloacae* showed a slower growth (∼1-2 log) in CFU at 24 h in Universal-Bac^3^Gel® relative to planktonic broth (purple vs blue bars, Figure **4A, 4J**), suggesting a transient adaptation phase. By 48 h, however, their counts in Universal-Bac^3^Gel^®^ had increased and were closer to planktonic levels (indicating these strains eventually proliferated well after the initial lag). In contrast, *A. baumannii* and *P. aeruginosa* maintained stable growth without any significant initial deficit. The number of bacteria in Universal-Bac^3^Gel® was similar to their planktonic counterparts at both 24 h and 48 h, indicating steady growth with no apparent adaptation lag (Figure **4D, 4G**).

**Figure 4.**
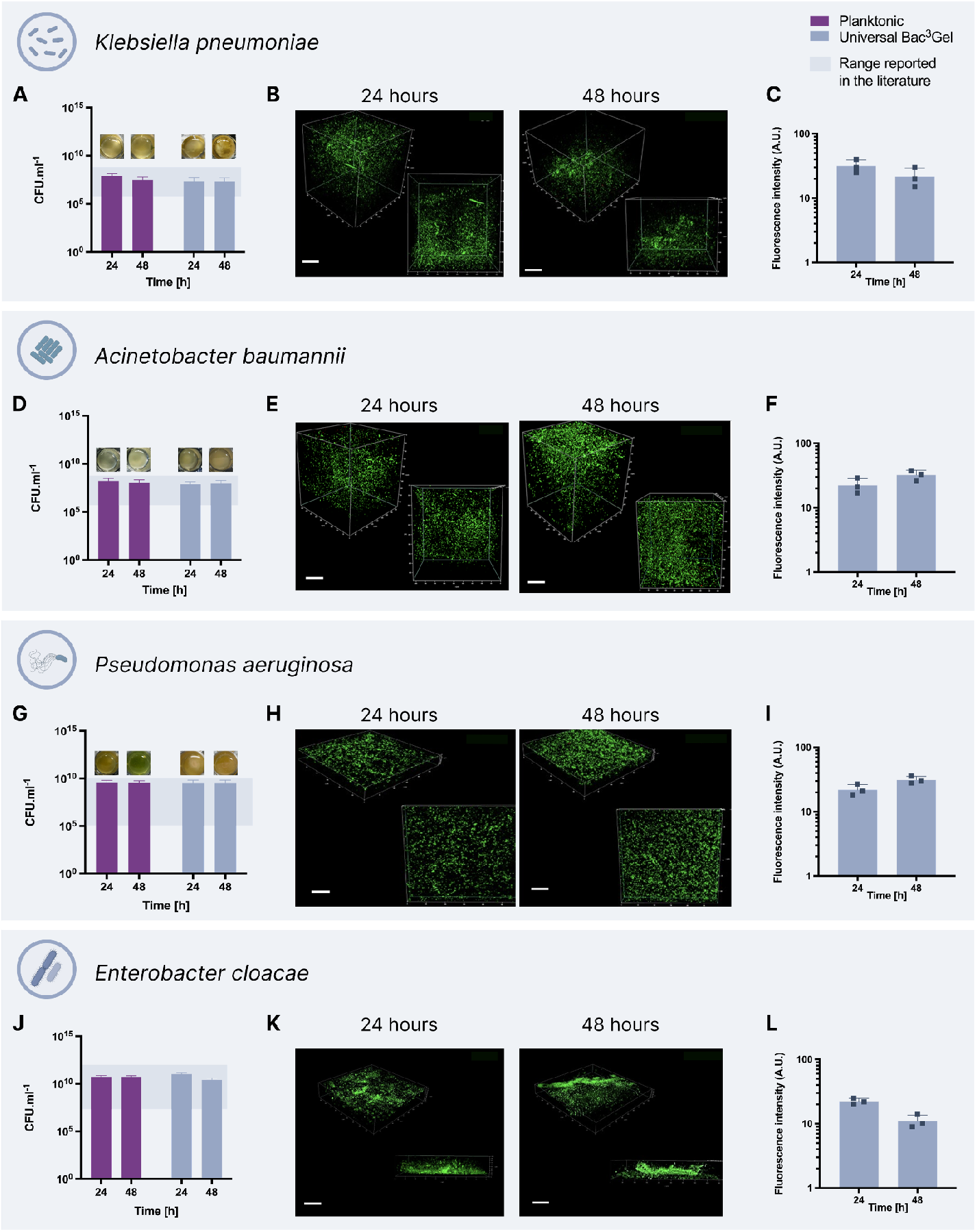
Growth and biofilm formation of Gram-negative ESKAPE pathogens in Universal-Bac^3^Gel®. (**A, D, G, J**) CFU counts of K. pneumoniae, A. baumannii, P. aeruginosa and E. cloacae at 24 h and 48 h in Universal-Bac^3^Gel® (purple) compared with planktonic cultures (blue. The shaded yellow area indicates the CFU/mL range reported for pathological mucosa. Data are mean ± SD (n = 3); p < 0.01 for all comparisons relative to 0 h inoculum. Photographs were taken only for the commercially available reference strains and not for clinical isolates. (**B, E, H, K**) Representative CLSM images showing live/dead staining in Universal-Bac^3^Gel®. Scale bar: 50 µm. (**C, F, I, L**) Quantification of green fluorescence intensity corresponding to viable biofilm biomass in 3D image stacks, illustrating total biofilm volume per field.

The intensity and distribution of green fluorescence signals in the 3D reconstructions correlate with viable bacterial biomass, supporting the ability of Universal-Bac^3^Gel® to sustain long-term colonization and species-specific biofilm architectures in Gram-negative pathogens. Microscopy-based qualitative analysis confirmed distinct 3D biofilm organization within Universal-Bac^3^Gel^®^ (**Figure 4B, 4E, 4H, 4K**). *K. pneumoniae* and *E. cloacae* formed compact yet dispersed aggregates, small clusters of cells scattered throughout the matrix, consistent with their capacity for early biofilm development. *A. baumannii* displayed a more homogeneous fluorescence throughout Universal-Bac^3^Gel^®^ (**Figure 4E**), suggestive of a diffuse colonization pattern with individual cells or small groups spread rather evenly. No large clusters were observed at 48 h observation after culturing *A. baumannii. P. aeruginosa*, by contrast, developed dense and spatially localized microcolonies (**Figure 4H**) - tightly packed spherical clusters - characteristic of its mature biofilm phenotype.

As with the Gram-positives, live/dead staining showed that the vast majority of cells in these biofilms were alive (green), with very limited red signals, indicating that Universal-Bac^3^Gel^®^ provides a conducive environment for the growth of these pathogens over 48 h. The intensity and distribution of green fluorescence correlated with viable bacterial biomass (**Figure 4C, 4F, 4I, 4L**).*P. aeruginosa* had very high localized green intensity (reflecting its large microcolonies), whereas *A. baumannii* showed a lower, evenly distributed signal. *K. pneumoniae* and *E. cloacae* had intermediate patterns. These species-specific biofilm architectures underscore the importance of using a 3D-relevant environment, as differences in spatial organization would be flattened or missed entirely in 2D culture.

## 4. Discussion

The results presented in this study demonstrate that Universal-Bac^3^Gel® provides a stable, reproducible, and physiologically relevant three-dimensional (3D) microenvironment for modeling biofilm-associated growth of ESKAPE pathogens (**Figure 1**). The hydrogel’s structural and rheological properties -namely its transparency, self-supporting nature, and viscoelastic behavior with a storage modulus (G′) higher than the loss modulus (G″) -closely resemble those of natural extracellular polymeric substance (EPS)-rich biofilm matrices (Klapper et al., 2002); (Pavlovsky et al., 2015). Indeed, the viscoelastic properties reported in the literature for natural biofilms -whose elastic moduli typically range from 0.2 to 25 Pa depending on species and maturation stage (Klapper et al., 2002); (Pavlovsky et al., 2015) closely match those of Universal-Bac^3^Gel®, whose storage modulus lies within the 2-200 Pa range (**Figure 2C-D**). The presence of a measurable oxygen gradient across the hydrogel depth further supports its ability to reproduce diffusion limitations typical of *in vivo* mucosal environments (**Figure 2B**). Together, these features generate a stratified habitat where microorganisms can proliferate and self-organize into biofilm-like communities.

A key outcome of this work is that all six ESKAPE pathogens maintained high viability and comparable CFU counts to their planktonic controls, confirming that Universal-Bac^3^Gel^®^ supports bacterial growth rather than imposing nutrient or diffusion constraints that limit proliferation. Importantly, confocal microscopy revealed pronounced differences in 3D spatial organization between species, showing that biofilm architecture within the hydrogel is both species-specific and influenced by the viscoelastic environment. These findings emphasize that the transition from planktonic to structured, multicellular organization is not solely dependent on surface attachment but can emerge spontaneously under mechanical confinement and diffusion gradients.

Among the Gram-positive pathogens, *E. faecium* exhibited a homogeneous distribution of viable cells forming a dense, lawn-like structure without large aggregates, consistent with a compact but dispersed biofilm phenotype (**Figure 3B**). This architecture suggests that *E. faecium* efficiently colonizes the 3D environment by uniform growth throughout the gel, possibly reflecting its adaptation to biofilm formation in nutrient-limited or fluid-rich tissues (Kim et al., 2021). In contrast, *S. aureus* formed spatially localized, dense aggregates typical of microcolony-based biofilms (**Figure 3E**). Such clustered architectures are hallmarks of *S. aureus* infection sites and have been associated with strong cell-cell adhesion, EPS accumulation, and oxygen gradient-driven stratification (Schilcher & Horswill, 2020). In Universal-Bac^3^Gel®, *S. aureus* exhibited more compact and localized aggregates throughout its matrix, compared to the previous gradient-based platform (Pacheco et al., 2023), possibly associated with the lack of mucin and the resulting reduction of biochemical adhesion cues.

For the Gram-negative species, distinct colonization patterns were also evident. *K. pneumoniae* and *E. cloacae* initially displayed slower growth at 24 h, suggesting a brief adaptation phase to the 3D environment, but by 48 h both species reached CFU levels comparable to planktonic cultures (**Figure 4A-J**). Microscopy analysis revealed compact, dispersed aggregates consistent with early-stage biofilm development and polymer-bridging aggregation mediated by exopolysaccharides (**Figure 4B-K**), a mechanism typical of these species in similar environments (Santajit & Indrawattana, 2016). *A. baumannii* (**Figure 4E**) showed a diffuse and homogeneous fluorescence pattern, indicative of dispersed colonization with limited aggregation. This morphology resembles the weakly cohesive, streamer-like biofilms formed by this pathogen under low-shear or nutrient-limited conditions, highlighting its tendency to form loose communities rather than structured clusters (Martí et al., 2011). In contrast, *P. aeruginosa* (**Figure 4H**) exhibited the most pronounced biofilm phenotype, developing dense, spherical microcolonies with intense fluorescence. These structures correspond to quorum-sensing-regulated EPS production and advanced biofilm maturation (Moradali et al., 2017) and mirror the stratified or layered architecture described in clinical mucus samples, where depletion aggregation and shear stress drive cellular stacking and co-aggregation (Savorana et al., 2025). Similarly to *S. aureus, P. aeruginosa* in Universal-Bac^3^Gel® exhibited a more homogeneous distribution pattern than in the previous gradient-based platform (Pacheco et al., 2023). This outcome further supports that mucin impacts bacterial spatial organization.

The diversity of architecture observed across species underscores how differences in bacterial physiology, extracellular matrix composition, and motility interact with the mechanical properties of the surrounding environment to determine biofilm morphology. Filamentous or clustered assemblies within Universal-Bac^3^Gel^®^ are consistent with recent physical models showing that cell shape and mechanical confinement regulate biofilm self-organization (Charlton et al., 2025). In such environments, filamentation lowers the percolation threshold and promotes the formation of connected networks through cluster-cluster aggregation and mechanical feedback. The viscoelastic properties of Universal-Bac^3^Gel^®^ likely provide similar mechanical cues: stiffness gradients, diffusion limitations, and stress relaxation collectively guide the emergence of anisotropic and filamentous growth, stabilizing biofilm structures over time. As highlighted by Savorana et al., 2025 the balance between storage and loss moduli (G′ > G″) determines the extent of mechanical coupling between cells and their surroundings, influencing biofilm cohesion, deformation, and recovery. The elastic modulus of Universal-Bac^3^Gel^®^ therefore not only supports physical stability but also promotes biologically meaningful organization and persistence.

From a biological perspective, structured 3D architectures confer several advantages to microbial communities, including enhanced antibiotic tolerance, metabolic cooperation, and protection from environmental stress (Pabst et al., 2016). The formation of species-specific morphotypes within Universal-Bac^3^Gel^®^ reflects adaptive strategies observed *in vivo*, such as microcolony formation by *S. aureus*, diffuse colonization by *A. baumannii*, and compact clustering by *K. pneumoniae*, demonstrating that the matrix effectively recapitulates key biofilm behaviors under controlled conditions. The combination of quantitative (CFU) and qualitative (CLSM) analyses confirms the robustness of the model in supporting both viability and spatial complexity, providing a reliable *in vitro* platform for studying microbial ecology, colonization dynamics, and biofilm physiology.

Overall, these findings establish Universal-Bac^3^Gel^®^ as a versatile and physiologically relevant system for investigating bacterial growth and organization in three dimensions. By enabling the formation of realistic biofilm architectures driven by viscoelastic confinement and diffusive gradients, the model bridges the gap between conventional planktonic assays and complex *in vivo* infection models. Its translational potential lies in providing a reproducible, ethically sustainable tool for a high throughput clinical testing of antimicrobials and anti-biofilm strategies, helping to better predict therapeutic efficacy against multidrug-resistant pathogens (Global Antibiotic Resistance Surveillance Report 2025 WHO Global Antimicrobial Resistance and Use Surveillance System (GLASS)

## 5. Conclusions

This study demonstrates that all six ESKAPE pathogens can successfully colonize and proliferate within the 3D Universal-Bac^3^Gel® matrix, a biomimetic hydrogel that replicates key structural and biochemical features of biofilm environments. This highlights its value as a relevant platform for studying biofilm formation and testing antimicrobial strategies under controlled conditions. By mimicking key aspects of the *in vivo* microenvironment, Universal-Bac^3^Gel® offers a promising tool for preclinical antimicrobial testing and could help bridge the gap between conventional 2D cultures and animal models. The inclusion of all ESKAPE priority pathogens in our evaluation underscores the broad applicability of the model. Future studies could leverage this system to test novel antibiotics, phage therapies, or anti-biofilm agents.

## Author Contributions

**Emanuela Peluso:**data curation (lead), formal analysis (lead), investigation (lead), methodology (equal), visualization (lead), writing–review and editing (lead). **Sebastião Van Uden:** conceptualization (lead), funding acquisition (lead), methodology (lead), project administration (supporting), resources (lead), validation (supporting). **Sonja Visentin:** conceptualization (supporting), funding acquisition (equal), project administration (supporting), supervision (supporting), validation (supporting). **Paola Petrini:** conceptualization (lead), funding acquisition (equal), project administration (supporting), supervision (supporting), validation (supporting). **Daniela Peneda Pacheco**: conceptualization (lead), funding acquisition (lead), methodology (lead), project administration (supporting), resources (lead), supervision (lead), validation (lead). **Livia Visai:** conceptualization (lead), funding acquisition (lead), project administration (lead), resources (lead) supervision (lead), validation (lead).

## Acknowledgments

The authors thank Amanda Oldani and Patrizia Vaghi (Centro Grandi Strumenti https://cgs.unipv.it/eng/, University of Pavia, Pavia, Italy) for technical assistance during CLSM analysis. L.V. and E.P. acknowledge the support from the Italian Ministry of University and Research (MUR) and the University of Pavia through the program “Dipartimenti di Eccellenza 2018–2022 and 2023-2027”. This project has been co-funded by the European Union’s Horizon Europe research and innovation programme under the European Innovation Council (EIC) Accelerator (HORIZON-EIC-2023-ACCELERATOROPEN-01 Grant Agreement No. 190135075) for the project Bac3Gel.

Created in BioRender. Visai, L. (2026) https://BioRender.com/bld7rwy

## Ethics Statement

None required.

## Data Availability Statement

The raw data supporting the findings of this study are not included in the article and are available from the corresponding authors upon reasonable request.

## Appendix 1. Additional Material

Link A1. Detailed protocol for Universal-Bac^3^Gel® assays.

Available at: https://drive.google.com/file/d/1Ln8UBYB1B8Pyxd_dd1LfK6v84iNwSX9v/view

## References

Aiyer, A., & Manos, J. (2022). The Use of Artificial Sputum Media to Enhance Investigation and Subsequent Treatment of Cystic Fibrosis Bacterial Infections. In Microorganisms (Vol. 10, Issue 7). MDPI. 10.3390/microorganisms10071269

Barrila, J., Crabbé, A., Yang, J., Franco, K., Nydam, S. D., Forsyth, R. J., Davis, R. R., Gangaraju, S., Mark Ott, C., Coyne, C. B., Bissell, M. J., & Nickerson, C. A. (2018). Modeling host-pathogen interactions in the context of the microenvironment: Three-dimensional cell culture comes of age. Infection and Immunity, 86(11). 10.1128/IAI.00282-18

Bermudez-Brito, M., Plaza-Díaz, J., Fontana, L., Muñoz-Quezada, S., & Gil, A. (2013). In vitro cell and tissue models for studying host-microbe interactions: A review. British Journal of Nutrition, 109(SUPPL. 2). 10.1017/S0007114512004023

Boucher, H. W., Talbot, G. H., Bradley, J. S., Edwards, J. E., Gilbert, D., Rice, L. B., Scheld, M., Spellberg, B., & Bartlett, J. (2009). Bad bugs, no drugs: No ESKAPE! An update from the Infectious Diseases Society of America. Clinical Infectious Diseases, 48(1), 1–12. 10.1086/595011

Bush, K., & Jacoby, G. A. (2010). Updated functional classification of β-lactamases. In Antimicrobial Agents and Chemotherapy (Vol. 54, Issue 3, pp. 969–976). 10.1128/AAC.01009-09

Butnarasu, C., Caron, G., Pacheco, D. P., Petrini, P., & Visentin, S. (2022). Cystic Fibrosis Mucus Model to Design More Efficient Drug Therapies. Molecular Pharmaceutics, 19(2), 520–531. 10.1021/acs.molpharmaceut.1c00644

Cone, R. A. (2009). Barrier properties of mucus. In Advanced Drug Delivery Reviews (Vol. 61, Issue 2, pp. 75–85). 10.1016/j.addr.2008.09.008

Crabbé, A., Jensen, P. ø., Bjarnsholt, T., & Coenye, T. (2019). Antimicrobial Tolerance and Metabolic Adaptations in Microbial Biofilms. In Trends in Microbiology (Vol. 27, Issue 10, pp. 850–863). Elsevier Ltd. 10.1016/j.tim.2019.05.003

Fasciano, A. C., Mecsas, J., & Isberg, R. R. (2019). New Age Strategies To Reconstruct Mucosal Tissue Colonization and Growth in Cell Culture Systems. Microbiology Spectrum, 7(2). 10.1128/microbiolspec.bai-0013-2019

Flemming, H. C., & Wuertz, S. (2019). Bacteria and archaea on Earth and their abundance in biofilms. Nature Reviews Microbiology, 17(4), 247–260. 10.1038/s41579-019-0158-9

Gilbert, J. A., Blaser, M. J., Caporaso, J. G., Jansson, J. K., Lynch, S. V., & Knight, R. (2018). Current understanding of the human microbiome. Nature Medicine, 24(4), 392–400. 10.1038/nm.4517

Global antibiotic resistance surveillance report 2025 WHO Global Antimicrobial Resistance and Use Surveillance System (GLASS). https://www.who.int/publications/i/item/9789240116337

Harrison, F., & Diggle, S. P. (2016). An ex vivo lung model to study bronchioles infected with Pseudomonas aeruginosa biofilms. Microbiology (United Kingdom), 162(10), 1755–1760. 10.1099/mic.0.000352

Higazy, D., Pham, A. D., Van Hasselt, C., Høiby, N., Jelsbak, L., Moser, C., & Ciofu, O. (2024). In vivo evolution of antimicrobial resistance in a biofilm model of Pseudomonas aeruginosa lung infection. ISME Journal, 18(1). 10.1093/ismejo/wrae036

Lieleg, O., & Ribbeck, K. (2011). Biological hydrogels as selective diffusion barriers. In Trends in Cell Biology (Vol. 21, Issue 9, pp. 543–551). 10.1016/j.tcb.2011.06.002

Möckel, M., Baldok, N., Walles, T., Hartig, R., Müller, A. J., Reichl, U., Genzel, Y., Walles, H., & Wiese-Rischke, C. (2022). Human 3D Airway Tissue Models for Real-Time Microscopy: Visualizing Respiratory Virus Spreading. Cells, 11(22). 10.3390/cells11223634

Mortensen, A. S., Nielsen, D. S., Røder, H. L., & Sølbeck Rasmussen, T. (2025). Mucofilm: a nexus for phage-microbiome interactions in gut ecology. Applied and Environmental Microbiology, 91(11). 10.1128/aem.01269-25

Munita, J. M., & Arias, C. A. (2016). Mechanisms of Antibiotic Resistance. Microbiology Spectrum, 4(2). 10.1128/microbiolspec.VMBF-0016-2015

Nasiri, R., Zhu, Y., & de Barros, N. R. (2024). Microfluidics and Organ-on-a-Chip for Disease Modeling and Drug Screening. In Biosensors (Vol. 14, Issue 2). Multidisciplinary Digital Publishing Institute (MDPI). 10.3390/bios14020086

Pacheco, D., Bertoglio, F., Butnarasu, C., Vargas, N. S., Guagliano, G., Ziccarelli, A., Briatico-Vangosa, F., Ruzzi, V., Buzzaccaro, S., Piazza, R., Uden, S. van, Crotti, E., Visentin, S., Visai, L., & Petrini, P. (2023). Heterogeneity governs 3D-cultures of clinically relevant microbial communities. 10.21203/rs.3.rs-2715275/v1

Pendleton, J. N., Gorman, S. P., & Gilmore, B. F. (2013). Clinical relevance of the ESKAPE pathogens. In Expert Review of Anti-Infective Therapy (Vol. 11, Issue 3, pp. 297–308). 10.1586/eri.13.12

Phogat, S., Thiam, F., Al Yazeedi, S., Abokor, F. A., & Osei, E. T. (2023). 3D in vitro hydrogel models to study the human lung extracellular matrix and fibroblast function. In Respiratory Research (Vol. 24, Issue 1). BioMed Central Ltd. 10.1186/s12931-023-02548-6

Ram, S., Lewis, L. A., & Rice, P. A. (2010). Infections of people with complement deficiencies and patients who have undergone splenectomy. In Clinical Microbiology Reviews (Vol. 23, Issue 4, pp. 740–780). 10.1128/CMR.00048-09

Rikken, G., Meesters, L. D., Jansen, P. A. M., Rodijk-Olthuis, D., van Vlijmen-Willems, I. M. J. J., Niehues, H., Smits, J. P. H., Oláh, P., Homey, B., Schalkwijk, J., Zeeuwen, P. L. J. M., & van den Bogaard, E. H. (2023). Novel methodologies for host-microbe interactions and microbiome-targeted therapeutics in 3D organotypic skin models. Microbiome, 11(1). 10.1186/s40168-023-01668-x

Santajit, S., & Indrawattana, N. (2016). Mechanisms of Antimicrobial Resistance in ESKAPE Pathogens. In BioMed Research International (Vol. 2016). Hindawi Limited. 10.1155/2016/2475067

Santos, J. de A., & Coelho, A. de A. (2025). Biopolymer Scaffolds in 3D Tissue Models: Advancing Antimicrobial Drug Discovery and Bacterial Pathogenesis Studies—A Scoping Review. Journal of Pharmaceutical and BioTech Industry, 2(3), 15. 10.3390/jpbi2030015

Sender, R., Fuchs, S., & Milo, R. (2016). Are We Really Vastly Outnumbered? Revisiting the Ratio of Bacterial to Host Cells in Humans. In Cell (Vol. 164, Issue 3, pp. 337–340). Cell Press. 10.1016/j.cell.2016.01.013

Shi, D., Mi, G., Wang, M., & Webster, T. J. (2019). In vitro and ex vivo systems at the forefront of infection modeling and drug discovery. Biomaterials, 198, 228–249. 10.1016/j.biomaterials.2018.10.030

Vargas, N. S., Antunes, M., Sobral, J., Silva, C., Sousa, F., Garbero, O. V., Kolková, A., Visai, L., Medana, C., Visentin, S., Petrini, P., van Uden, S., & Pacheco, D. (2025). Gut 3 Gel: A High Throughput Mucus Model for Culturing Human Intestinal Microbiota. 10.1101/2025.02.21.639490

Zeden, M. S., & Gröndling, A. (2023). Agar Plate-Based Method for the Selection of Antibiotic-Resistant Bacterial Strains. Cold Spring Harbor Protocols, 2023(11), 816–821. 10.1101/pdb.prot107897

